# Physiological and ecological warnings that Dodder pose an exigent threat to farmlands in Eastern Africa

**DOI:** 10.1101/2020.10.26.355883

**Authors:** Joel Masanga, Beatrice Njoki Mwangi, Willy Kibet, Philip Sagero, Mark Wamalwa, Richard Oduor, Mathew Ngugi, Amos Alakonya, Patroba Ojola, Emily S. Bellis, Steven Runo

**Affiliations:** Department of Biochemistry, Microbiology and Biotechnology. Kenyatta University, Kenya; Kenya Meteorological Department, Nairobi, Kenya; International Maize and Wheat Improvement Center, Mexico; Arkansas Biosciences Institute and Department of Computer Science, Arkansas State University, USA; Center for No-Boundary Thinking, Arkansas, USA

**Keywords:** *Cuscuta*, *Cuscuta* resistance, Parasitic plants, species distribution models

## Abstract

Invasive holoparasitic plants of the genus *Cuscuta* (dodder) threaten Africa’s ecosystems, due to their rapid spread and attack on various host plant species. Most *Cuscuta* species cannot photosynthesize, hence rely on host plants for nourishment. After attachment through a peg-like organ called a haustorium, the parasites deprive hosts of water and nutrients leading to their death. Despite their rapid spread in Africa, dodders have attracted limited research attention, although data on their taxonomy, host range and epidemiology are critical for their management. Here, we combine taxonomy and phylogenetics to reveal presence of field dodder (*Cuscuta campestris*) and *C. kilimanjari* (both either naturalized or endemic to East Africa), and for the first time in continental Africa, presence of the giant dodder (*C. reflexa*) a south Asian species. These parasites have a wide host range, parasitizing species across 13 angiosperm orders. Evaluating the possibility of *C. reflexa* to expand this host range to tea, coffee, and mango, crops of economic importance to Africa, revealed successful parasitism, following haustorial formation and vascular bundle connections in all three crops. However, only mango mounted a successful post-attachment resistance response. Furthermore, species distribution models predicted high habitat suitability for all three *Cuscuta* species across major tea- and coffee-growing regions of Eastern Africa, suggesting an imminent risk to these crops. Our findings provide relevant insights into a little-understood threat to biodiversity and economic wellbeing in Eastern Africa, and providing critical information to guide development of management strategies to avert their spread.

**Sentence Summary:** Microscopy and habitat suitability modeling provide an early warning that dodder’s invasion in Eastern Africa poses a threat to important cash crops

## Introduction

Invasive parasitic plants are a major threat to plant communities, due to their profound negative impacts on global biodiversity and agricultural productivity (Press and Phoenix, 2005). In Africa, some parasitic plants, such as *Striga spp.* are well researched because of their direct negative impacts on cereal staples (reviewed by Parker, 2012). However, others such as dodder (*Cuscuta spp*.), noxious vines of the Convolvulaceae family, have received little attention.

The genus *Cuscuta* comprises over 200 species of obligate parasites that infect a wide range of herbaceous and woody plants, including important crop species (Lanini and Kogan, 2005). *Cuscuta spp*. are widely distributed across the world, and reportedly colonize a wide range of hosts across various habitats (Lanini and Kogan, 2005). Overall, members of this genus occur on all continents, except Antarctica, with most species reported in the Americas and Mexico, which are also considered their centre of diversity (Yuncker, 1932; Stefanović *et al*., 2007). In Africa, only a handful of studies have reported dodder occurrence (Zerman and Saghir, 1995; Garcia, 1999; Garcia and Martin, 2007; Garcia *et al*., 2014). However, their distribution patterns remain unknown. According to the Flora of Tropical East Africa (Verdcourt, 1963), several species, namely *C. australis*, *C. campestris* Yuncker, *C. suaveolens* Seringe, *C. kilimanjari* Oliv, *C. hyalina* Roth, *C.* Engelm, *C. epilinum* and *C. planiflora* Tenore, are endemic to or naturalized in Eastern Africa. However, some non-native species have recently been anecdotally reported although information about where they were introduced from is unclear.

Cuscuta’s widespread success is attributed to its parasitic life history strategy and ability to steal most of the resources needed for growth and reproduction from its hosts. Particularly, most *Cuscuta* species do not photosynthesize, due to reduced levels (or lack thereof) of chlorophylls, although some show localized photosynthesis (Parker and Riches, 1993; Braukmann *et al*., 2013; Kim and Westwood, 2015). Consequently, they entirely depend on their hosts for nourishment.

*Cuscuta* life cycle follows a systematic pattern that begins with seed germination, attachment and penetration of a suitable host (through a specialized organ called haustorium), development of vegetative tissues, flowering and seed production. Due to a limited amount of food reserves in their seeds, seedlings must attach to an appropriate host within 3-5 days of germination (Lanini and Kogan, 2005), and establish vascular bundle connections that act as a conduit for siphoning water, nutrients and photo-assimilates. Thereafter, the parasite develops flowers and eventually produces viable seeds that shed back to the soil (Dawson *et al.*, 1994; Albert *et al.*, 2008). At this point, the host succumbs to parasitism often leading to its death.

Morphological identification of *Cuscuta* species is difficult because of lack of morphological descriptors based on leaf structre. All members have slender vines, with scale-like leaves and no roots. Previous taxonomic identification relied on floral and fruit characters. In this regard, the early monograph by Engelmann, (1857) categorized *Cuscuta* into 3 groups, based on stigma and style morphology. These were later adopted by Yuncker, (1932) as subgenera. Specifically, subgenus Monogynella is characterized by fused styles, whereas subgenera Cuscuta and Grammica have 2 distinct styles, distinguished by respective elongate and globose stigmas. Yuncker later revised the monograph and subdivided these subgenera into 8 sections, based on fruit dehiscence, and 29 subsections based on a combination of characters, such as flower numbers, size, texture and shape, as well as density of inflorescence among others.

We hypothesized that different *Cuscuta* species currently occur in Eastern Africa, and their distribution patterns are due to various biotic factors, such as presence of suitable hosts, and interactions with the environment. This is because, in general, occurrence of a species in a particular locality is shaped by life history characteristics, environmental requirements, population genetics and their associations with ecology over time. Therefore, we first used a combination of morphological descriptors and sequencing of the plastid locus (Ribulose bisphosphate carboxylase large- *rbcL* and *trnL*) as well as nuclear ITS region, to identify *Cuscuta* species presently invading ecosystems in Kenya. We then determined their host range by compiling a comprehensive list of current *Cuscuta* hosts and extrapolated the possibility of the parasite to expand this range to crop trees, by infecting coffee (*Coffea arabica*), tea (*Camelia sinensis*) and mango (*Mangifera indica*) under screenhouse conditions. We selected these crops because of their agricultural/economic importance. In Kenya, they contribute to the country’s GDP through export earnings and cover an estimated area of 114,700, 218,538 and 60,497 ha for coffee, tea and mango respectively (FAO, 2017).

Finally, we used geographical information system (GIS)-based species distribution modelling (SDM) to estimate geographical distribution of the identified *Cuscuta spp*. across Eastern Africa, based on current climatic conditions and vegetation. Specifically, we adopted presence-only SDMs using the maximum entropy (MaxEnt) algorithm, which combines occurrence records with environmental variables to build correlative models for predicting habitat suitability for a species (Phillips *et al*., 2006). This algorithm has been previously used to predict distribution of parasitic plants, such as *C. chinensis* (Ren *et al*., 2020), *Striga hermonthica* (Cotter *et al*., 2012) and, mistletoes (Zhang *et al*. 2016).

We found that the current dodder invasion in Kenya i) comprises *C. campestris*, *C. kilimanjari* and *C. reflexa*; ii) has a wide host range that could potentially include tea and coffee; and iii) has a wide distribution with potential to invade new habitats. These findings will inform policies for management of the parasite in Eastern Africa.

## Results

### Floral morphological characters reveal 3 *Cuscuta* species

Floral characters revealed 3 distinct *Cuscuta* species in accessions collected from Kenya (Fig. 1). Summarily, flowers across all specimens had clusters comprising 5 petals, 5 sepals and 5 stamens, and were identified to the species level as follows:

***C. campestris* Yuncker (subgenus Grammica)**: accessions here comprised slender, threadlike yellow to orange stems with a diameter of about 0.3 mm (Supplemental Fig. S1A). Flowers were small, and white, about 2 mm in diameter, with greenish-yellow capsules that appeared in compact cymose clusters. Calyx lobes were obtuse, or somewhat acute, whereas corolla lobes were triangular. Stamens were shorter than the lobes, with filaments of about 1mm. They had 2 separate slender styles, about 1 mm long, with globose stigmas that did not split at the base. Four ovules, about 1 mm long, were present (Fig. 1Aa-d).
***C. kilimanjari* Oliv**: accessions here had thick, coarse and purple vines, about 1 mm in diameter (Supplemental Fig. S1B). Flowers were pale white, waxy, about 4 mm wide. Both sets of calyx and corolla were obtuse, whereas stamens were shorter than the lobes, with short and thick filaments. This category had 2 separate short and thick styles, less than 1 mm long and 0.3 mm wide. Styles bore white spherical stigmas, with ovaries that had purple spots (Fig. 1Ba-d).
***C. reflexa* Roxb**: vines were greenish-yellow, >2 mm in diameter (Supplemental Fig. S1C), with large flowers about 6 mm wide. The flowers had a single thick and short style, with 2 elongated stigmas, and ovaries that contained 4 white ovules, of different sizes (Fig. 1Ca-d).

**Fig. 1.**
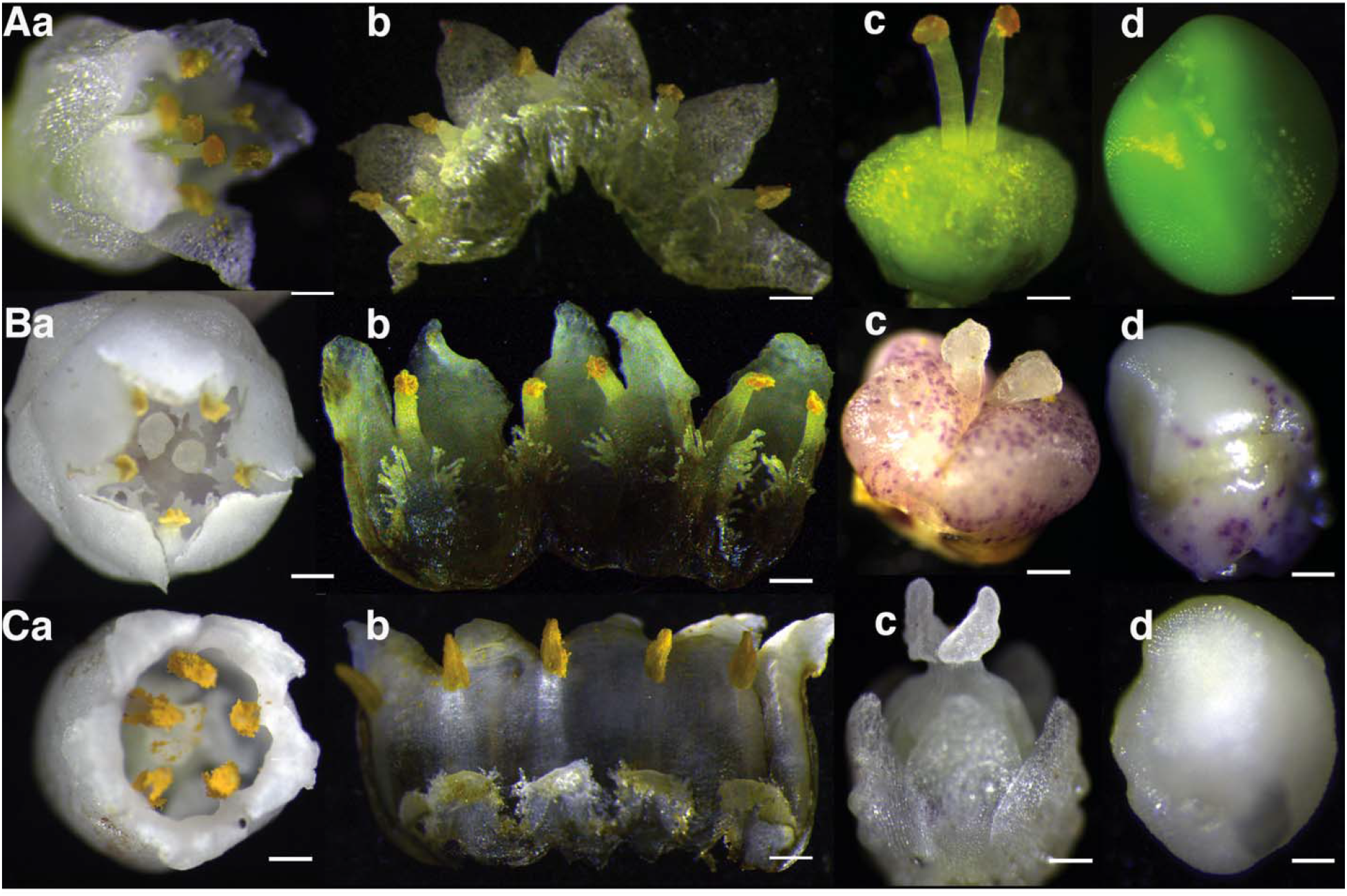
Profiles of floral morphology among *Cuscuta* accessions collected across Kenya, showing variations in gynoecia, ovule shape, size and colour across species. Aa-Ad *C. campestris*- evidenced by small, white flowers with separate styles that bear globose stigmas; Ba-Bd *C. kilimanjari*- confirmed by thick separate styles with spherical stigmas; Ca-Cd *C. reflexa*- evidenced by short fused styles that bear ligulate stigmas. Bars Aa = 0.4 mm; Ab = 0.4 mm; Ac = 0.4 mm; Ad = 0.1 mm Ba = 0.8 mm; Bb = 0.8 mm; Bc = 1 mm; Bd = 1 mm; Ca = 1.2 mm; Cb = 1.2 mm; Cc = 1 mm and Cd = 0.2 mm.

### Phylogenetic analysis

Parsimony consensus trees for accessions under this study, alongside other *Cuscuta* species from GenBank, revealed consistent topologies across all 3 (*rbcL, trnL* and ITS) regions (Fig. 2). In brief, 3 major clades, corresponding to the 3 *Cuscuta* subgenera, were resolved across the datasets with good bootstrap support. Specifically, *C. campestris* accessions from the current study were resolved in the first major clade with other reported species of subgenus Grammica, sister to *C. campestris* taxa from GenBank (Fig. 2). Within the same clade, our *C. kilimanjari* accessions were resolved and nested with a South American clade of subgenus Grammica. The second major clade comprised members of subgenus *Cuscuta*, with emphasis on species previously reported to occur in Africa. The third major clade comprised subgenus Monogynella, with our *C. reflexa* taxa nested inside a Genbank-derived *C. reflexa* group and basal to both subgenera *Cuscuta* and Grammica. Unrooted Maximum Likelihood gene trees confirmed that subgenus Monogynella were basal to subgenus Grammica, across all genes tested, as well as in the combined dataset (Supplemental Fig. S2)

**Fig. 2.**
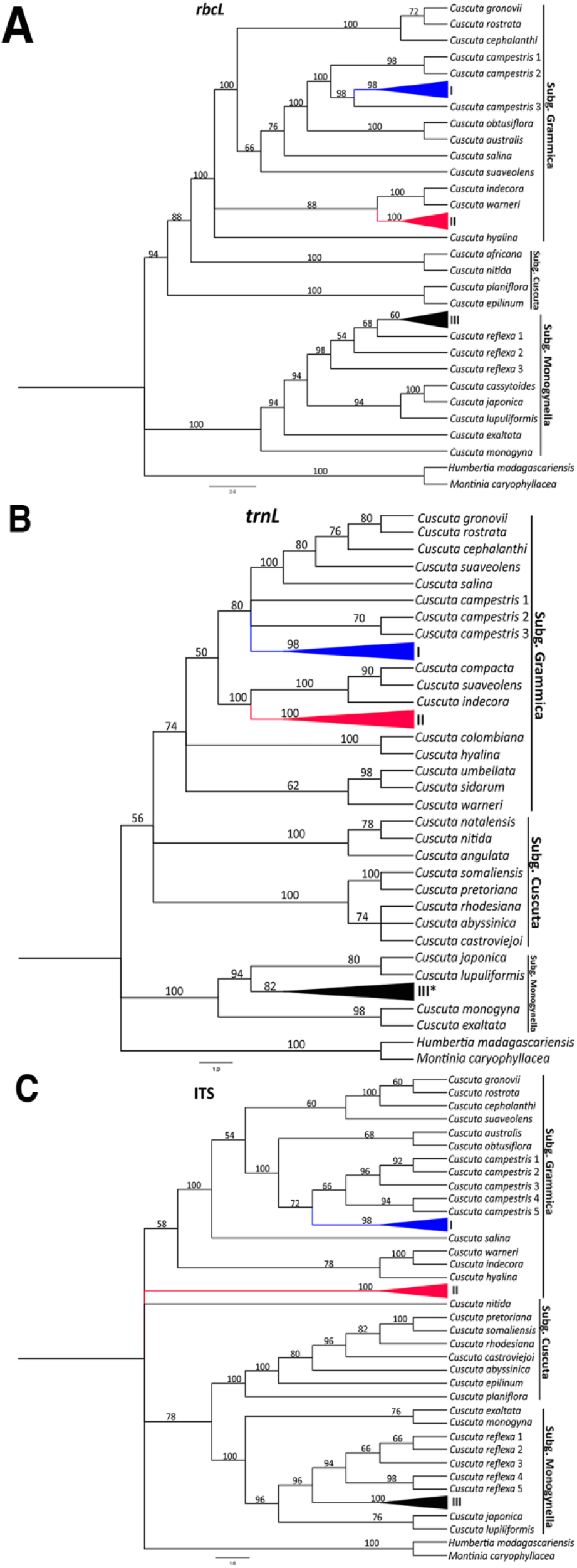
Phylogenetic reconstruction of *Cuscuta* species based on *rbcL*, *trnL* and ITS regions. Maximum Parsimony bootstrap consensus trees (1000 replicates) are shown, with bootstrap supports indicated above branches. I, II and III represent *Cuscuta* taxa sequenced under this study, and denote *C. campestris*, *C. kilimanjari* and *C. reflexa*, respectively. The Asterisk (*) on the *trnL* tree implies that our *C. reflexa* taxa were collapsed with those from GenBank.

### Dodder has a wide host range with potential to infect crops of great economic importance

A total of twenty-six (26) host plant species across 13 angiosperm orders, comprising shrubs (40%), trees (44%) and herbs (16%) were parasitized by the aforementioned *Cuscuta* species. Fabales (20%) was the most parasitized order, followed by Lamiales (16%), Malpighiales and Caryophyllales (both with 12%), whereas the rest had a single species colonized by the parasite.

With regards to host specificity, *C. campestris* and *C. reflexa* exhibited a ‘generalist’ behaviour, indiscriminately parasitizing hosts across numerous orders. Among the parasitized species, *Solanum incanum* and *Biancaea decapetala* were the most preferred hosts for *C. campestris*, whereas *Thevetia peruviana* was the most preferred host for *C. reflexa*. Conversely, *C. kilimanjari* exhibited a ‘specialist’ behaviour, parasitizing host species across 2 orders only (Supplemental Table. S1).

We further demonstrated the parasite’s potential threat to crops by infecting tea, coffee and mango with *C. reflexa,* followed by histological analysis. We focused on *C. reflexa* for infections due to its invasiveness across the region, a wide host preference (perennial trees and shrubs) and because it was introduced to E. Africa. In all instances (100%), *Cuscuta* successfully parasitized test plants and formed haustoria within 14 days of infection. Cross sections, performed 4 weeks after infection, revealed successful penetration of the parasite into host tissues, past the cortex and endodermis, enabling successful formation of vascular bundle connections (Fig. 3). Interestingly, we observed a resistance response from an infected mango plant. Specifically, the infected point swelled, and exuded a sap-like substance that was deposited around the wounded area. This eventually led to death of the parasite (within 4 weeks of attachment), with the infected area ‘healing’ afterwards (Supplemental Fig. S3).

**Fig. 3.**
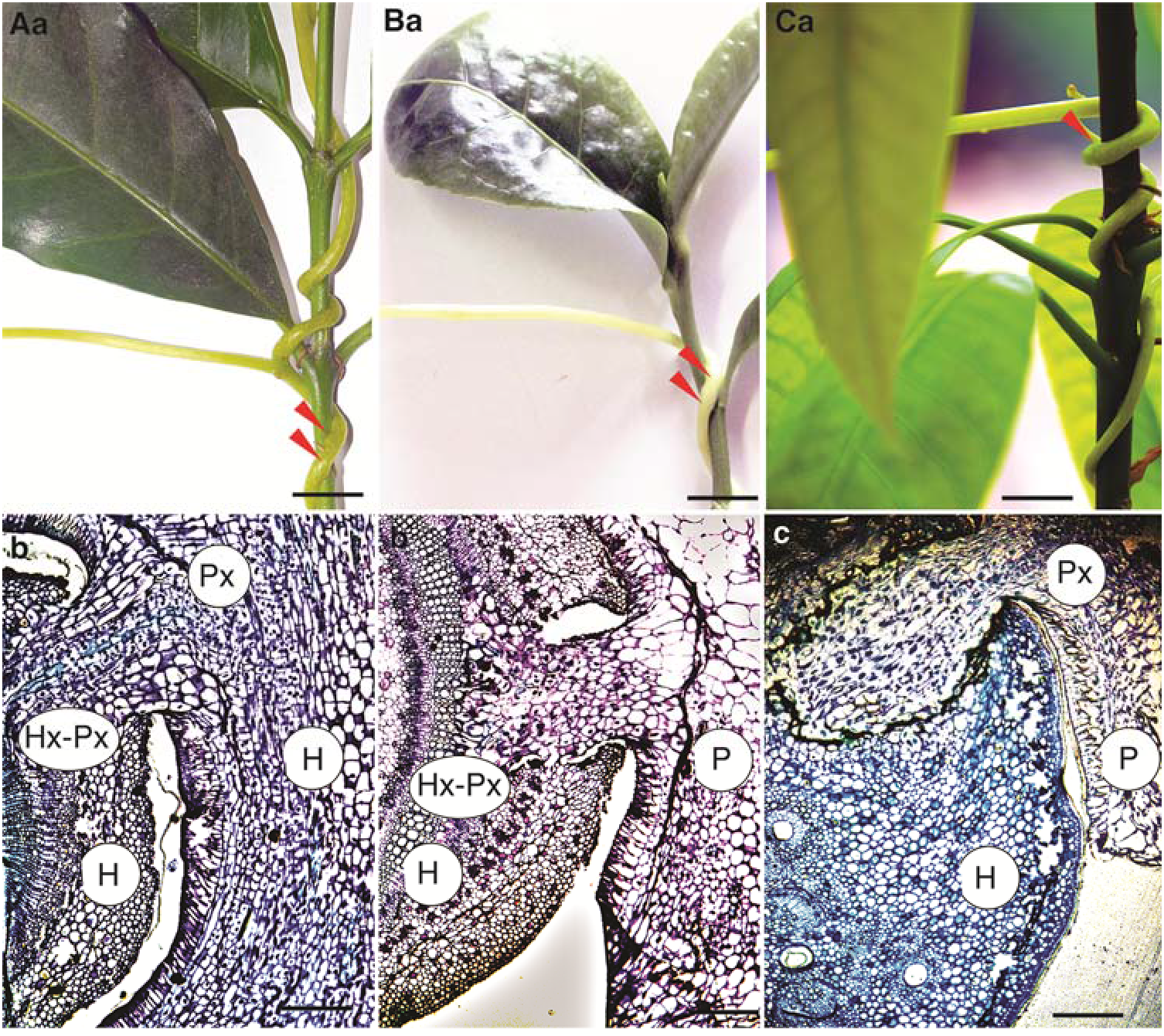
*Cuscuta* parasitism and extent of ingression into host plants. The upper panel shows close up photographs of infected test plants while the lower panel are toluidine blue-stained cross sections of the host-parasite interface. Aa and Ab- coffee, Ba and Bb-tea, Ca and Cb- mango. P-parasite; H-host; HX-Host xylem; PX-Parasite xylem. Bar top panel=10 mm, bottom panel=5 mm.

To further evaluate the imminent danger posed by these parasites to tea, we sampled Kenyan locations where the ranges for tea and *C. reflexa* overlapped. As an example, we highlight a site in Kakamega, Western Kenya. https://earth.google.com/web/search/0+12%272%27%27N,+34+46%2721E/@0.202444,34.77314623,1583.10168457a,0d,15y,119.88850412h,90.54106391t,0r/data=CigiJgokCdGS14h74TRAEc6S14h74TTAGbGLz-dnrjzAIbWQBbGXbGDAIhoKFnVHbklCUWhkQ1BPdnMyZll4TTFqdFEQAg

Here, we found *Markhamia lutea*, a host of *C. reflexa* was infected and growing just next to a tea plantation – pointing to the definite possibility of tea infestation. A Google Earth™ image (using the Street View option) taken in June 2018 showed that the tree had not been infested, but by the time we visited the site in August 2019, the tree had heavy infestation that threatened to encroach the tea plantation (Fig. 4). This indicates that *C. reflexa* is highly invasive with potential to rapidly infest new localities. In its native ranges of Asia, *C. reflexa* has been reported to parasitize a wide range of hosts, including coffee (Bhattarai *et al*., 1989; Das, 2007). Additionally, dodder has been reported on coffee in Uganda (Jennipher Bisikwa Personal Communication), and one of the records at the East African Herbarium (voucher number EA16731, collected in 1983) indicated that one of the *C. kilimanjari* specimens parasitized coffee.

**Fig. 4.**
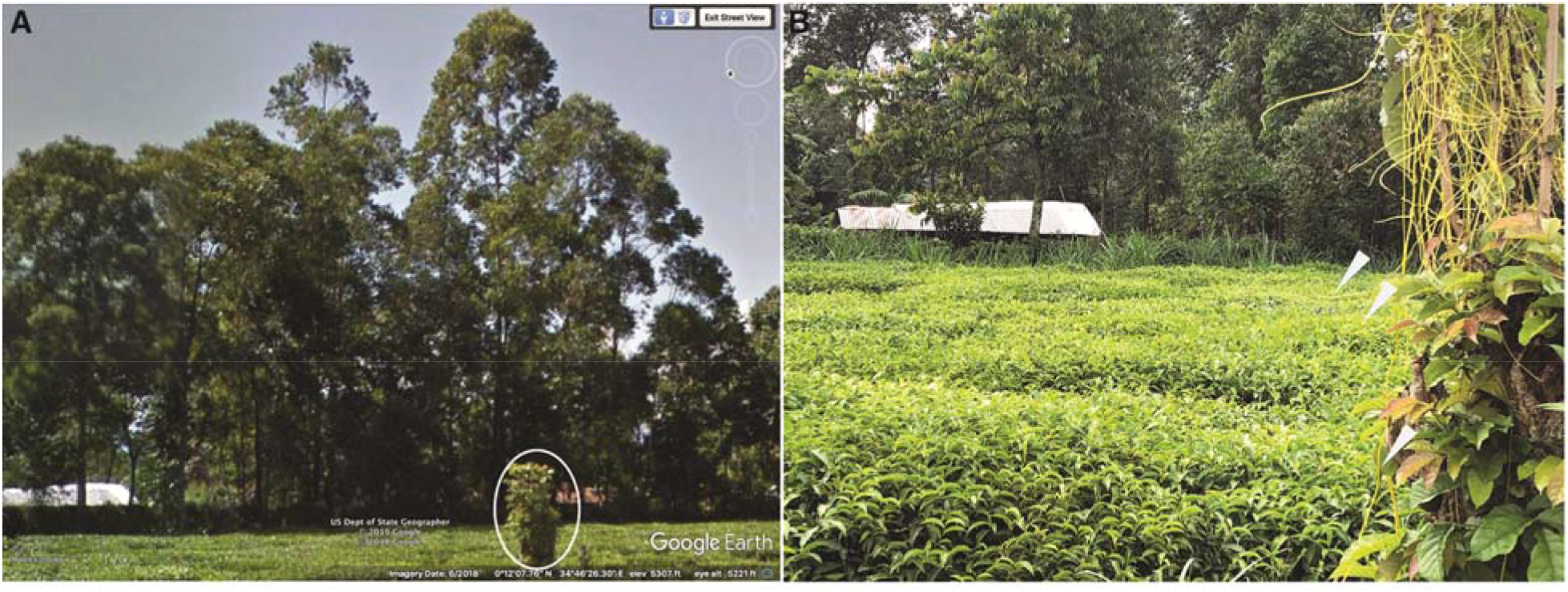
*Cuscuta* threat on tea. **A** Google Earth™ (Street View) image of a tea plantation in western Kenya, taken in 2018. The circled bush represents *M. lutea*, a tree species commonly used as a windbreaker around plantations and later found to be a *Cuscuta* host. **B** The windbreaker infested with *C. reflexa,* one year after the first image. Arrows indicate parasitic vines threatening to encroach into the tea plantation.

### Predicted *Cuscuta* distribution and habitat suitability

Our SDMs had excellent predictive performances, with AUC values of 0.93 and 0.87 for *C. reflexa* built using occurrence records from Kenya and native ranges, respectively. On the other hand, models for *C. campestris* and *C. kilimanjari* had modest performances, with AUC values of 0.76 and 0.72, respectively (Supplemental Figs S4-S7). Precipitation of the warmest quarter (bio18=59.0%) and annual mean temperature (bio1=32.2%) were the most influential variables in the models for *C. reflexa* based on occurrences in Kenya and the native range, respectively. Conversely, precipitation of the driest quarter (bio17=46.2) and isothermality (bio3=47.6%) were the highest contributors to the models for *C. campestris* and *C. kilimanjari*, respectively (Table 1).

**Table 1.** Permutation importance of bioclimatic and vegetation variables. Values represent percent (%) contributions of each variable to the model. An asterisk (*) in each column indicates the highest contributing variable

| **Variable** | ***C. campestris*** | ***C. kilimanjari*** | ***C. reflexa* (invaded)** | ***C. reflexa* (native)** |
| --- | --- | --- | --- | --- |
| Annual Mean Temperature (bio1) | 4.4 | 0 | 0 | 32.2* |
| Isothermality (bio3) | 2.5 | 47.6* | 13.7 | 1.1 |
| Precipitation Seasonality (bio15) | 0 | 12.7 | 0.5 | 15.6 |
| Precipitation of Driest Quarter (bio17) | 46.2* | 2.7 | 0.5 | 15.9 |
| Precipitation of Warmest Quarter (bio18) | 4.9 | 7.7 | 59.0* | 13.2 |
| Land Cover Fraction (Grass) (veg1) | 40.1 | 25.8 | 14.9 | 1.7 |
| Land Cover Fraction (Shrub) (veg2) | 0.3 | 0 | 0.8 | 11.2 |
| Land Cover Fraction (Tree) (veg3) | 1.5 | 3.5 | 10.6 | 9.1 |

Our models presented in Fig. 5 revealed different distribution patterns across Eastern Africa, with current estimates showing that all 3 species under this study can potentially colonize areas larger than the localities sampled herein.

**Fig. 5.**
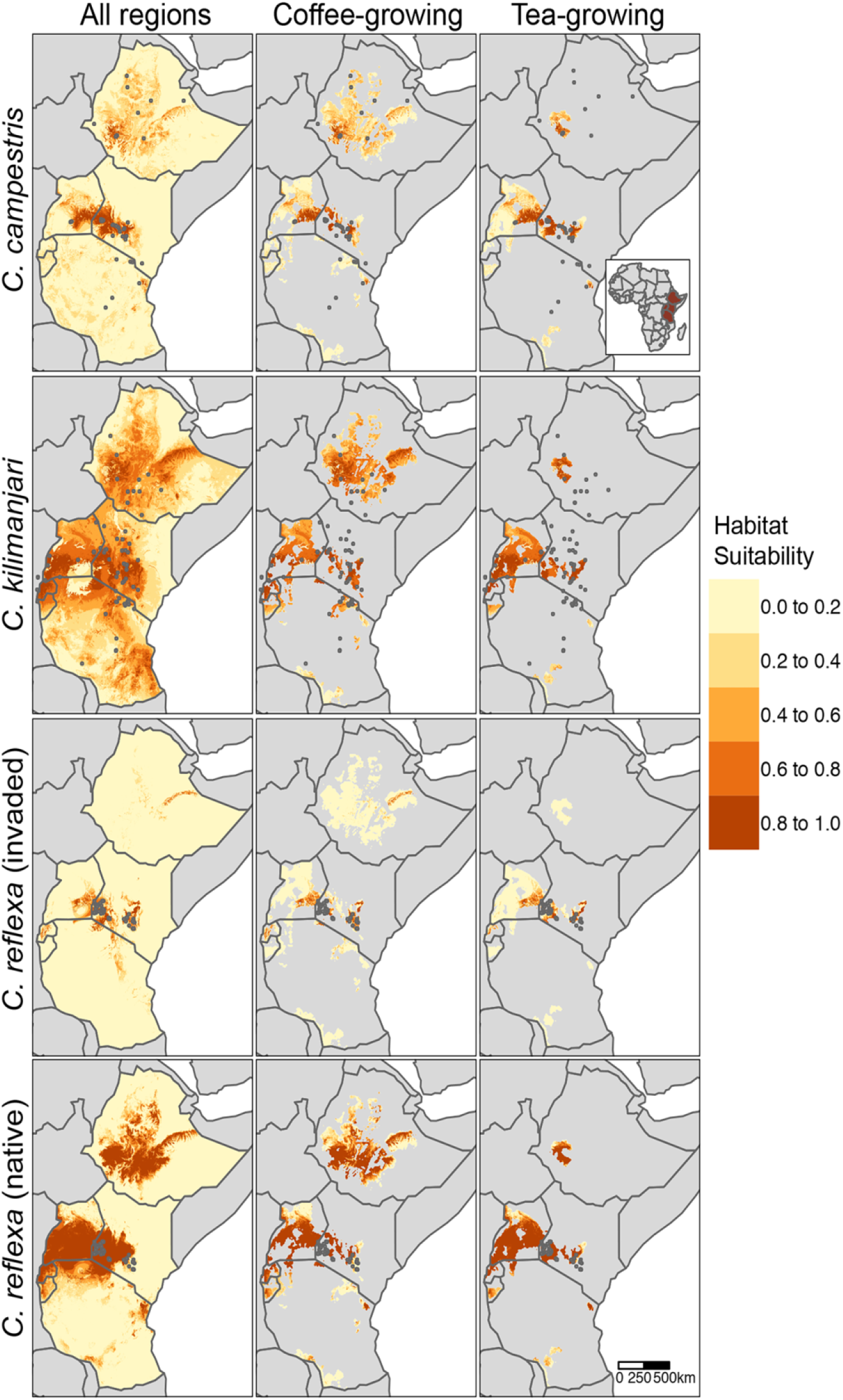
Habitat suitability for *C. campestris, C. kilimanjari, C. reflexa* across Ethiopia, Kenya, Uganda, Rwanda, Burundi, and Tanzania. Dark grey points indicate locations of occurrence records. For *C. reflexa*, species distribution models were trained using occurrences from the invaded range in Kenya (*n*=66) or native range from Afghanistan to Indo-China (*n*=165), then projected to the six countries. Models for *C. campestris* and *C. kilimanjari* were constructed using a combination of occurrence records obtained from our sampling activities (in Kenya), as well as those obtained from GBIF and herbarium specimens at the East African Herbarium. Projections were masked to coffee- and tea-growing regions, estimated to produce >1 metric tons in 2017 (IFPR, 2020).

Overall, high habitat suitability for *C. reflexa* (log-transformed score >0.8) is predicted in Western and Central Kenya, Eastern Uganda, large parts of Rwanda and Burundi as well as Central Ethiopia. Predicted habitat suitability for *C. kilimanjari* is higher than that observed for *C. reflexa* across the 6 countries. Particularly, Central and Western Kenya, Western Uganda, Rwanda and Burundi, as well as Central Ethiopia show high suitability. Conversely, moderate habitat suitability is predicted for *C. campestris*, with infestation likely to occur in Western and Central Kenya, Eastern Uganda, with pockets in Ethiopia and Tanzania. Additionally, many major coffee- and tea-growing areas show high habitat suitability for all species. Our collected occurrence data indicate that *C. reflexa* seems to have already invaded many of these regions in Kenya. Projections of our models to other regions suggest that *C. reflexa* could become or already may be a concern in coffee- and tea-growing regions of Rwanda, eastern Uganda, and the Harar coffee zone of Ethiopia.

Noteworthy, the predicted distributions are supported by various local press describing dodder infestations in the aforementioned countries including Uganda (https://www.newvision.co.ug/new_vision/news/1503872/dangerous-plant-invades-kampala-city) and Kenya (https://www.nation.co.ke/news/Dodder-plant-poses-threat-to-trees-and-crops/1056-5138904-aawhphz/index.html).

## Discussion

### *Cuscuta* identification

Interactions among global change components, such as land use, agricultural intensification and invasions by non-native species, cause substantial changes to plant community composition worldwide. Consequently, plant communities that arguably play the biggest role in establishing and maintaining ecosystems are deteriorating. A key driver of such community change is invasion by alien plants that displace/arrest development of native species, subsequently reducing habitat quality, causing economic losses, and posing a serious threat to human wellbeing (Vila’ *et al*., 2011). Once established, the spread of invasive plants is extremely difficult to control or reverse. In African terrestrial ecosystems, a myriad of invasive plant species occurs, key among them the current rapid spread of parasitic vines of the genus *Cuscuta*. To avert loss to biodiversity and developed reasoned management programs for the parasites, we analysed *Cuscuta*’s potential threat in Eastern Africa by describing their taxonomy, host range and habitat suitability.

*Cuscuta* identification using morphological characters is difficult because members of this genus lack roots and their leaves are reduced to minute scales (Kuijt, 1969). However, diversity among floral components presents a unique opportunity for distinguishing species. We adopted monographs by Engelmann (1857) and Yuncker (1932) to interpret stigma and style morphology, and consequently identified 3 *Cuscuta* species, from 2 related subgenera (Grammica and Monogynella) in accessions collected from Kenya. Individuals in subgenus Grammica had 2 separate styles, with either globose or spherical stigmas, and were classified as *C. campestris* and *C. kilimanjari*, respectively, whereas those in subgenus Monogynella had elongate stigmas born on fused styles, typical of *C. reflexa*.

We validated this identification by sequencing *rbcL, trnL* and ITS regions from representative individuals, and performed phylogenetic reconstruction alongside other species from GenBank. These markers have been extensively used to infer phylogenetic relationships, character evolution and biogeography across the genus (Garcia and Martin, 2007; McNeal., *et al*., 2007; Garcia *et al*., 2014). In our case, all our sequences were resolved alongside respective *C. campestris, C. kilimanjari* and *C. reflexa* taxa from GenBank, indicating that they indeed belonged to these species. Additionally, these phylogenetic reconstructions confirmed the monophyly of subgenus Monogynella, with all members basal to subgenera *Cuscuta* and Grammica, consistent with earlier reports (Garcia and Martin, 2007; McNeal *et al*., 2007; Stefanović *et al*., 2007; Garcia *et al*., 2014). Apart from these 3, other *Cuscuta* species have also been reported in Africa, although most of them belong to subgenus *Cuscuta* (Zerman and Saghi, 1995; Garcia, 1999; Garcia and Martin, 2007).

### SDM-based distribution patterns

*Cuscuta*’s damaging success and worldwide distribution is attributed to its ability to colonize a wide range of hosts and survive in areas with an array of environmental conditions (Parker and Riches, 1993). Additionally, most members are non-specific, colonizing multiple host plants across various angiosperm families (Dawson *et al*., 1994; Lanini and Kogan, 2005; Kim and Westwood, 2015).

Our SDMs, combined with the observed host range and artificial infection assays, provided insights into the parasite’s potential distribution patterns in Eastern Africa. These models showed that the species are likely to be present in additional areas not covered by this study, as evidenced by areas of high habitat suitability.

Three climatic variables, precipitation of the warmest quarter, annual mean temperature and isothermality had the highest contribution to our models, suggesting that they could play a significant role in the predicted distribution. These were also found to significantly contribute to habitat suitability for *C. chinensis* (Ren *et al*., 2020).

With regards to land cover contribution, grass cover fraction was an important predictor for SDMs in all three species, with lower habitat suitability corresponding with areas of high grass cover. This finding is consistent with lower frequency of preferred *Cuscuta* hosts in grasslands and parasitism on diverse herbaceous plants as well as perennial shrubs and trees.

Additional identification of lower risk areas for *Cuscuta* invasion is critical for future preparedness. For example, our models predicted relatively low habitat suitability in central and western Uganda for *C. reflexa* (according to models based on occurrences from a restricted sampling area in the invaded range). However, SDMs based on occurrences throughout a broader set of environments in the native range suggested high habitat suitability in these same regions. Although there are many limits to spatial transferability of SDMs from native and introduced ranges (Liu *et al*., 2020), these findings suggest that current environments may not be an effective barrier to the spread and establishment of *C. reflexa* in many East Africa regions and that active management of *C. reflexa* will be needed to prevent its spread.

### *Cuscuta*’s potential threat to cash crops

Results from artificial infection of *C. reflexa* on coffee, tea and mango revealed its potential to parasitize these crops of economic importance. Specifically, we observed attachment, haustoria and vascular bundle formation; all indicative of successful parasitism. *Cuscuta* parasitism on coffee and tea could have devastating impacts on the income generated by these crops in East African countries. In addition, presence of a *C. reflexa*-infected *M. lutea* just next to a tea plantation is indicative that such an infestation may be imminent. Governments in East Africa will therefore require to develop urgent interventions and appropriate policies to stop such eventualities which would have devastating effects to farmers in affected areas.

Interestingly, we observed a resistance response in the mango genotype artificially infected with *C. reflexa*. Cross sections indicated ingression of parasitic haustoria into the host and successful establishment of vascular bundle connections. However, this success was short-lived with the host initiating wound response and chemical deposition that resulted in death of the parasite and later, healing of the infected area. It is possible that this resistance response goes beyond the observed wounding, although this remains to be investigated. Such a phenomenon could also be key in determining dodder’s host preference since plants that display resistance are avoided during parasitism (Kaiser *et al.*, 2015). Previous studies have described this type of resistance (Albert *et al*., 2006) among other mechanisms, including incompatibility due to anatomical attributes (Dawson *et al.*, 1994), induction of defence-related stress hormones (salicylic and jasmonic acid) as well as use of mechanical barriers that block parasitic ingression into host vasculature (Kaiser *et al.*, 2015). Consequently, species such as *Gossypium hirsutum* (Capderon *et al*., 1985), *Solanum lycopersicum* (Albert *et al*., 2006; Runyon *et al.*, 2010) and some varieties of chickpea (*Cicer arietinum*) (Goldwasser *et al.*, 2012) have been reported to resist dodder infection and are therefore avoided during host selection.

### Conclusions and future prospects

In summary, our findings reveal presence of 3 *Cuscuta* species; *C. campestris, C. kilimanjari* and *C. reflexa* across various ecosystems in Kenya. The first 2, endemic to or naturalized in Eastern Africa, have been documented in the Flora of Tropical East Africa, alongside others such as *C. australis*, *C. suaveolens* Seringe, *C. hyalina* Roth, *C. cassytoides* Engelm, *C. epilinum* and *C. planiflora* Tenore (Verdcourt, 1963). However, to the best of our knowledge, this is the first report describing occurrence of *C. reflexa*, a south Asian species, in continental Africa. These parasites have a broad host range, and infestation on crops of economic importance may be inevitable if urgent actions are not taken to stop their spread. In fact, our predictions show that many regions across Eastern Africa are characterized by highly suitable habitats for *Cuscuta* infestation, and may already be infested. Therefore, this work will be critical in developing informed strategies for managing the parasite and averting the looming risk. This may potentially involve identifying resistant plant species and genotypes to aid development of cultural control and adaptation measures in agriculture and forestry within the region. Additionally, unravelling the physical, biochemical and genetic factors controlling the observed resistance response in mango will provide insights into regulation of these resistance phenomena and guide future control strategies.

## Materials and methods

### Sample collection

We collected a total of 96 *Cuscuta* accessions across Kenya. In our case, an accession is defined as material from an individual plant from the same species found within a similar geographical area. Sampling was done between July and November 2018, with at least 5 individual accessions collected per location. Dodder flowers and vines were collected and immediately dried in silica gel to await morphological analysis and DNA isolation. *Cuscuta*-parasitized plants were also collected and identified to the species level, according to the keys of plant identification described in the Flora of Tropical East Africa (Verdcourt, 1963) and Pennsylvania State University (https://extension.psu.edu/plant-identification-preparing-samples-and-using-keys).

### Morphological *Cuscuta* identification

Morphological identification was performed according to the keys of *Cuscuta* monograph constructed by Engelmann, (1857) and Yuncker, (1932). Briefly, flowers were either used immediately after collection or rehydrated before microscopy (for those kept in silica gel). To observe different parts, we examined a single full flower (sepals, petals, gynoecium and androecium) under a Leica MZ10F stereomicroscope (Leica Microsystems, UK) and photographed it. Thereafter, we carefully dissected and photographed it, with focus given to the gynoecia, number of parts, fusion (or lack thereof) of the styles as well as the size and shape of stigmas. Ovaries were also dissected, then the number, size and color of ovules observed and photographed.

### Host plant infection and histology

We evaluated whether dodder could expand its host range to tree crops of agricultural value, by artificially infecting tea, coffee and mango with *C. reflexa,* under controlled conditions in the greenhouse. Summarily, three months-old test seedlings were maintained in potted soil with regular watering then infected by winding a 30 cm long piece of parasitic vine (that had at least one node) around their stems. Parasitism was determined by histological analysis of the host-parasite interface, four weeks after infection as previously described (Gurney *et al.*, 2003).

Briefly, tissues at the interface were collected and fixed using Carnoy’s fixative (4:1 ethanol: acetic acid), dehydrated with 100% ethanol then pre-infiltrated in Technovit solution (Haraeus Kulzer, GmbH). The tissues were embedded in 1.5 ml microcentrifuge tube lids containing Technovit/Hardener and left to set according to the manufacturer’s instructions, then mounted onto histoblocks using the Technovit 3040 kit (Haraeus Kulzer GmbH). Microscopic sections (5 μm thick) of the tissues were cut using a microtome (Leica RM 2145), transferred onto glass slides and stained using 0.1% Toluidine Blue O dye in phosphate buffer. After washing off excess dye and drying, slides were covered with slips containing a drop of DPX (BDH, Poole, UK), observed and photographed using a Leica microscope mounted with a Leica MC190 HD camera (Leica, UK).

### DNA extraction, polymerase chain reaction (PCR) and Sequencing

We sequenced representative accessions from each of the aforementioned *Cuscuta* species, following morphological characterization. We could not acquire material for DNA extraction from voucher specimens held at the East African Herbarium in Nairobi, Kenya, owing to the destructive nature of sampling involved. DNA was extracted from flowers and hanging vines, collected at least 10 cm away from the point of attachment to avoid host-DNA contamination.

PCR amplification of the ITS region was done using ITS4 and ITS5 primers (Baldwin, 1992), whereas *rbcL* was amplified using *rbcL*-512F and *rbcL*-1392R primers (McNeal *et al*., 2007). Partial amplification of *trnL* was using *trnL*F-5’ CGAAATCGGTAGACGCTACG 3’ and *trnL*R-5’ ATTTGAACTGGTGACACGAG 3’ primers, designed specifically for *Cuscuta*. PCR reactions were performed in 25 μl volumes using MyTaq™ DNA polymerase kit (Bioline, Meridian Biosciences) under the following conditions; 95°C for 1 minute, followed by 35 cycles comprising 95°C for 15 seconds, each primer’s respective annealing temperature for 30 seconds and a 72°C extension for 1 minute. A final 10-minute extension, at 72°C, was also included. PCR products were confirmed on a 1% agarose gel, cleaned using the Qiaquick™ PCR purification kit (Qiagen, USA), and sequenced on the ABI platform at Macrogen (Macrogen. Inc).

### Phylogenetic analysis

Sequences were edited in SeqMan Pro17 in Lasergene package (DNASTAR Inc., Madison, WI, USA) to remove low quality reads, then aligned using ClustaX version 2.0 (Larkin *et al*., 2007). Sequences were submitted to NCBI (Accession numbers are shown at the end of the document), then used as queries to identify similar taxa using the nucleotide BLAST algorithm at NCBI. Highly similar sequences across the 3 *Cuscuta* subgenera were retrieved for phylogenetic reconstruction and ancestry inferencing, with sequences for *Montinia caryophyllacea* and *Humbertia madagascariensis* included as outgroups. We first constructed unrooted Maximum Likelihood gene trees for the accessions under this study, then generated Parsimony consensus trees (with 1000 replications) for phylogenetic reconstruction of taxa from our species alongside those from NCBI. All phylogenetic analyses were performed in MEGAX (Kumar *et al*., 2018), and the trees visualized in FigTree version 1.4.4 (Rambaut, 2009).

### Species distribution modelling

#### Occurrence records

We combined respective occurrence records for *C. kilimanjari* and *C. campestris* from our sampling efforts in Kenya with records from the Global Biodiversity Information Facility (GBIF) (https://doi.org/10.15468/dl.9kpzum), and records for specimens held in the collection at the East African Herbarium (EA). This resulted in a total of 74 and 51 unique locations for *C. kilimanjari* and *C. campestris*, respectively. We found no records of *C. reflexa* in neither GBIF nor in the EA collection, hence all occurrence records for models based on this species’ current invasion range were from localities sampled as part of this study (*n* = 66 unique locations across Kenya). We also built SDMs based on 165 unique occurrences of *C. reflexa* from its native range (Afghanistan to Indo-China) that were available from GBIF (https://www.gbif.org/occurrence/download/0074148-200613084148143). To characterize the background of the study, we randomly sampled 1,000 points from a radius of 300 km from known occurrences.

#### Environmental variables

Species distribution models were based on five bioclimatic variables, namely annual mean temperature (bio1), isothermality (bio3), precipitation seasonality (bio15), precipitation of the driest quarter (bio17), and precipitation of the warmest quarter (bio18), as well as three variables related to vegetation structure, namely land cover fraction of grass (veg1), shrub (veg2), and tree (veg3). Four of the bioclimatic variables (bio1, bio3, bio15, and bio18) were previously reported as important for species distribution models for *C. chinensis* (Ren *et al*., 2020), whereas the fifth (bio17) exhibited high feature importance in our preliminary analyses. Bioclimatic data were obtained from the CHELSA dataset (https://zenodo.org/record/3939050#.X3N49y2ZM8Z) (Karger *et al*., 2017), whereas vegetation layers were from the Copernicus Global Land Service (Buchhorn *et al*., 2020). These layers were based on epoch 2019 from Collection 3 of the annual, global 100m land cover maps, and were resampled to match resolution of the bioclimatic layers (1 km) using the bilinear interpolation method of the ‘resample’ function from the R package ‘raster’ (Hijmans 2019). The vegetation layers captured aspects of vegetation cover, which may be important for *Cuscuta* spp. parasitism on various herbaceous and woody host species. All variables had Pearson’s correlation coefficients less than 0.8 across background points of the study.

### Model building and prediction of suitable habitats

Species distribution models were built using the Maxent algorithm (Phillips *et al*., 2006). The Models were tuned and evaluated with R version 3.6.1 with ENMeval (Muscarella *et al*., 2014) using the checkerboard2 method for partitioning occurrence data into training and test sets. To determine overlap between *Cuscuta* spp. distributions with major coffee- and tea-growing areas, we used crop production maps from IFPRI, (2020) (https://dataverse.harvard.edu/dataset.xhtml?persistentId=doi:10.7910/DVN/FSSKBW) to mask *Cuscuta* spp. distribution models to just those areas estimated to produce at least one metric ton of coffee or tea in 2017.

## Accession Numbers

***<u>rbcL</u>***

***C. campestris***; MW078922, MW078923, MW078924

***C. kilimanjari***; MW078930, MW078931, MW078932

***C. reflexa***; MW078927, MW078928, MW078929

**ITS**

***C. campestris***; MT947605, MT947606, MT947607

***C. kilimanjari***; MT952140, MT952141, MT952142

***C. reflexa;*** MW080817, MW080818, MW080819

***trnL***

**C. campestris**; MW086603, MW086604, MW086605

***C. kilimanjari***; MW086607, MW086608, MW086609

***C. reflexa***; MW115588, MW115589, MW115590

## Supplemental Data

The following supplemental materials are available

**Fig. S1** Categories of *Cuscuta* species observed parasitizing various susceptible host plants in Kenya.

**Fig. S2** Unrooted Maximum Likelihood trees based on *rbcL, trnL*, ITS and a combination of the 3 regions.

**Fig. S3** Resistance response exhibited by a mango (*Mangifera indica*) genotype under *C. reflexa* infection.

**Fig. S4** Area under curve (AUC) values for *C. campestris*.

**Fig. S5** Area under curve (AUC) values for *C. kilimanjari.*

**Fig. S6** Area under curve (AUC) values for *C. reflexa*

**Fig. S7** Area under curve (AUC) values for *C. reflexa*

**Supplemental Table S1** Occurrence records of *Cuscuta* species collected from Kenya

## Acknowledgements

We thank the National Museums of Kenya, through the East Africa Herbarium, for providing occurrence records. We acknowledge Prof. Claude dePamphilis (Pennsylvania State University-USA) for fruitful scientific discussions and Prof. Alistair Jump (University of Stirling-UK) for thoughtful reviews.

